# Bigger is not always better: size-dependent fitness effects of adult crowding in *Drosophila melanogaster*

**DOI:** 10.1101/2025.04.21.649761

**Authors:** Medha Rao, Chinmay Temura, Avani Mital, S. Anvitha, Amitabh Joshi

## Abstract

Density-dependent selection is an important factor shaping the evolution of life histories. In holometabolous insects, crowding in the larval and adult stages can have very different effects on key fitness components. While the nuanced effects of density-dependent selection through larval crowding in *Drosophila melanogaster* have been extensively studied for various life history traits, very few studies have investigated the effects of adult crowding in *Drosophila*. Moreover, these few studies were mostly conducted on large flies, derived from low larval density cultures, and typically treated the overall density of flies per culture container as an index of the strength of adult crowding. We hypothesized that the size of the adults should shape the impact of adult crowding, with small individuals experiencing less stress than large individuals when crowded. Consequently, the adverse fitness effects usually associated with adult crowding may not be observed for small individuals. We tested this hypothesis by subjecting flies of different sizes – regular-sized flies, and small flies derived via larval crowding or selection for rapid development to adulthood – to an episode of adult crowding and examining their mortality and fecundity. Thus, we explored the interactive effects between larval and adult crowding on key fitness components. Small body size enabled flies to handle adult crowding better, with significantly lower mortality under crowded conditions when compared to flies of large body size. Moreover, small flies showed a consistent pattern of increased fecundity upon adult crowding. This positive impact on fecundity was not observed when larger flies were crowded. It is clear from our study that the effects of adult crowding can be very nuanced and body size-specific, even to the extent of having a net beneficial effect on fitness components, contrary to previous belief.

## Introduction

The survival and reproduction of organisms is often affected by the limited availability of resources, such as food, space and mates (Mueller, 1988; Andersson, 1994). The life-history of organisms can thus be shaped, both ecologically and evolutionarily, by the density they experience with regard to a specific limiting resource (Mueller & Joshi, 2000; Prasad & Joshi, 2003). This notion forms the foundation of the theory of density-dependent selection, based on the idea that the relative fitness of a variant may be density-specific (reviewed in Mueller, 1997; Travis *et al*., 2023). In many species, especially of holometabolous insects such as *Drosophila*, larval and adult life-stages can have very different form and ecology (Demerec & Kaufmann, 1996). Consequently, the selection pressures experienced in each life-stage can also differ in kind and intensity. In this scenario, selection experienced at one life-stage can significantly impact subsequent life-stages (Mueller, 1988; Chippindale *et al*., 1997; Prasad & Joshi, 2003). For example, the amount of food consumed by a *Drosophila* larva has been shown to affect subsequent adult body size and female fecundity (Robertson & Sang, 1944a,b). Selection for rapid egg-to-adult development, which largely acts on the larval stage in *Drosophila*, is known to have significant effects on the adult stage, with adults exhibiting a significantly lower body weight at eclosion (Nunney, 1996; Chippindale *et al*., 1997; Prasad *et al*., 2000), along with lower female fecundity (a correlate of body size at eclosion; Prasad, 2003) and reduced longevity (Modak, 2009).

While density-dependent selection in *Drosophila* has been studied in considerable detail (reviewed in Mueller, 1986; Prasad & Joshi, 2003), most work focused on the effects of crowding experienced during the larval stage and examined the impact of larval crowding on larval and adult fitness components (Bakker, 1962; Mueller *et al*., 1991; reviewed in Mueller, 1997). A common implicit assumption of these studies was that the number of larvae present per unit volume of food medium (overall density) was a reliable index of the strength of larval competition. The canonical view of the evolutionary consequences of high larval density that emerged from these studies was one in which the principal adaptations that evolved in populations experiencing chronic larval crowding were an increase in larval feeding rate (Joshi & Mueller, 1988, 1996) and tolerance to metabolic waste (Shiotsugu *et al*., 1997; Borash *et al*., 1998), which was achieved in part at the cost of efficiency of food conversion to biomass (Joshi & Mueller, 1996; Mueller, 1990), contrary to the previously held view that crowding would favour the evolution of greater food to biomass conversion efficiency (MacArthur & Wilson, 1967; Mueller, 1988).

However, subsequent studies challenged both this canonical view and the notion that overall larval density is a good index of the strength of larval competition. One set of studies showed that the outcomes of larval density-dependent selection in *Drosophila* could depend critically on the details of how exactly (the precise combination of egg number and food volume) crowding was imposed on the larvae, and that greater larval competitive ability could evolve without a concomitant increase in larval feeding rate (Nagarajan *et al*., 2016; Sarangi *et al*., 2016; Sarangi, 2018). In a follow-up study examining the ecological consequences of larval crowding using different numbers of eggs placed at different combinations of food column height and surface area, it was found that the effective density (number of larvae per unit volume of food present in the feeding band) was a better predictor of the outcome of crowding than the overall density of larvae per unit volume of food (Venkitachalam *et al*., 2023). Thus, it was recognized that differences in the ecological details of how larval crowding was enforced, even at the same overall density, could greatly affect the outcome of crowding on various fitness-related traits such as body size, egg-to-adult development time and pre-adult survivorship (Venkitachalam, 2023).

Compared to larval crowding, the ecological and evolutionary effects of adult crowding in *Drosophila* have been relatively less studied (reviewed in Prasad & Joshi, 2003). A study (Pearl & Parker, 1922) found that fertility (the number of adult progeny produced per female) significantly decreased when flies were subjected to increased adult density in bottle cultures. Subsequent studies (Pearl, 1927, 1932) observed increased mortality at high adult densities and showed that female fecundity in *Drosophila* decreased with increasing adult density. They found the optimal adult density for a population to be around 35-55 flies per bottle, with mortality rates rising on either side of this threshold. Another study (Robertson & Sang, 1944a) noted that the surface area available for oviposition, combined with the nutritional status of the flies, governed patterns of fecundity upon experiencing adult crowding. A further study (Mueller & Huynh, 1994) examined the effects of adult crowding in combination with the nutritional quality of food (with or without supplemental yeast-acetic acid paste) on the fecundity of females. They found that the fecundity of females decreased with an increase in adult density. The decrease in fecundity was very sharp, especially when the food was not supplemented with yeast. Across studies, it was clear that a relatively short duration (24 hours) of adult crowding was enough to significantly affect adult mortality, fertility, and the fecundity of females.

In addition to the few studies examining the immediate fitness effects of adult crowding summarized above, there has been one experimental evolution study that contrasted adaptation to adult crowding experienced every generation with adaptation to chronic larval crowding. This study involved three sets of 5 replicate populations each, selected for adaptation to crowding, either at the larval (CU populations) or the adult stages (UC populations), and their controls (UU populations) (first described in (Mueller *et al*., 1993). These populations were examined for their fitness responses to adult crowding by examining longevity, fecundity and mortality after experiencing adult crowding (Joshi, 1997; Joshi & Mueller, 1997; Joshi *et al*., 1998). The UC populations had evolved better adaptations to adult crowding, with lower mortality on exposure to adult crowding and higher fecundity of the females after an episode of adult crowding, relative to the UUs and CUs. For each type of population, a consistent pattern of decreased fecundity was observed upon experiencing a short period of adult crowding (Joshi *et al*., 1998).

These studies gave rise to the notion that the number of adults within a given space is a reliable indicator of the intensity of adult competition. The consensus formed from the studies was that adult crowding resulted in a net negative effect on components of fitness like mortality and fecundity. However, some of the early studies used inbred lines of *Drosophila* for their assays, limiting the generality of their results (Pearl, 1932; Robertson & Sang, 1944a). Interestingly, a common feature across all these studies was that flies used for the assay were reared at low larval density. Low larval density entails rearing larvae obtained from a small number of eggs in a high volume of food. The density at which larvae were reared for the assays (∼30-80 eggs in ∼4-8 mL of food across studies) would ensure that the emerging adults from such cultures would be large, with very low variation in body size. Therefore, while the range of tested adult crowding densities varied, the flies used were consistently large due to the similar nature of larval rearing.

We hypothesized that the body size of the individuals influences the effects of adult crowding on key fitness components. We hypothesized that small flies would be less affected by adult crowding compared to larger flies. Compared to the large flies used in earlier studies, small flies could perhaps better deal with the space limitation created by adult crowding. If the space limitation experienced is less severe, the effect of adult crowding might not be as detrimental as seen in earlier studies and could even have a positive effect. In this context, we decided to explore the response of smaller-sized adults to adult crowding in contrast to larger adults.

Flies of smaller body sizes used in our assay were obtained using different methods, as elaborated below. Selection for various traits can result in a correlated response of body size. Studies have shown that populations of *Drosophila* that have been selected for rapid development and early reproduction (FEJs), as well as populations that have been relaxed to various degrees from the above selection pressures (CRFs, FRFs), have evolved a significantly smaller body size relative to their ancestral controls (JBs) as a correlated response to selection (Mital, 2019). Another method to obtain smaller body size is through larval crowding, which tends to give rise to adults of significantly smaller size (Lints & Lints, 1969; Scheiring *et al*., 1984). If the small flies obtained by different methods showed a similar response to adult crowding, we could conclude that the response was likely due to the body size, not the method by which the size was obtained.

Our experimental design allowed us to examine the interactive effects of imposing both larval and adult crowding in populations of *D. melanogaster*, and we asked if flies with small body sizes, achieved through different means, converged in their response to adult crowding. We exposed flies to a short duration (48 h) of adult conditioning density (low or high) and measured the effect of the adult density in two ways – the mortality incurred during the adult conditioning and the fecundity of surviving females.

Our results indicate that female fecundity decreased as adult density increased for flies reared at a low larval density (large flies), consistent with results from earlier studies. However, for flies reared at high larval density (small flies), an increase in adult density resulted in a corresponding rise in female fecundity. Our study is the first report of a positive impact of adult crowding on fecundity, a key fitness component. It is also the first to show that the body size of the individuals experiencing crowded conditions plays a role in shaping the net outcome of adult crowding.

## Results

### Effect of age, body size and evolutionary history on fecundity

Fecundity on day 12 was ∼27.4% higher than on day 18, indicating a significant impact of the age of the flies on fecundity (Table 1, main effect of day, *F*_1, 3_ = 19.54, *p* = 0.021). Flies reared at low larval density (L-LD) developed into heavier adults than those reared at high larval density (H-LD; Table S1, main effect of larval density, *F*_1, 3_ = 226.172, *p* = 0.042). Larger flies (reared at L-LD) had higher fecundity than smaller flies (reared at H-LD) (Table 1, main effect of larval density, *F*_1, 3_ = 221.1, *p* <0.001).

**Table 1.**
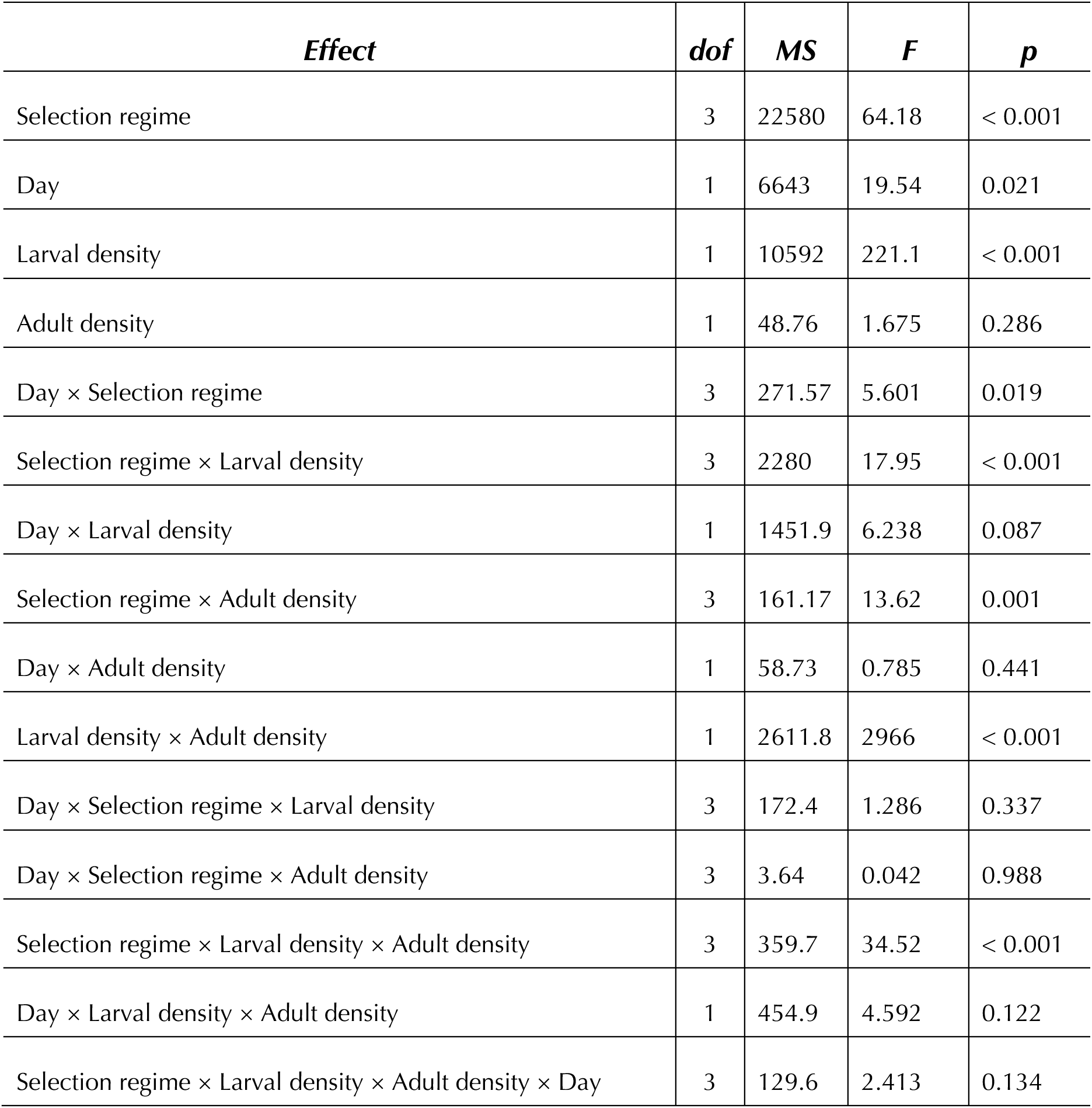
ANOVA results for fecundity of female flies after the adult conditioning period. In this design, random factors and their interactions are not tested for significance and are omitted from the table.

| *Effect* | *dof* | *MS* | *F* | *p* |
| --- | --- | --- | --- | --- |
| Selection regime | 3 | 22580 | 64.18 | < 0.001 |
| Day | 1 | 6643 | 19.54 | 0.021 |
| Larval density | 1 | 10592 | 221.1 | < 0.001 |
| Adult density | 1 | 48.76 | 1.675 | 0.286 |
| Day × Selection regime | 3 | 271.57 | 5.601 | 0.019 |
| Selection regime × Larval density | 3 | 2280 | 17.95 | < 0.001 |
| Day × Larval density | 1 | 1451.9 | 6.238 | 0.087 |
| Selection regime × Adult density | 3 | 161.17 | 13.62 | 0.001 |
| Day × Adult density | 1 | 58.73 | 0.785 | 0.441 |
| Larval density × Adult density | 1 | 2611.8 | 2966 | < 0.001 |
| Day × Selection regime × Larval density | 3 | 172.4 | 1.286 | 0.337 |
| Day × Selection regime × Adult density | 3 | 3.64 | 0.042 | 0.988 |
| Selection regime × Larval density × Adult density | 3 | 359.7 | 34.52 | < 0.001 |
| Day × Larval density × Adult density | 1 | 454.9 | 4.592 | 0.122 |
| Selection regime × Larval density × Adult density × Day | 3 | 129.6 | 2.413 | 0.134 |

Among the selection regimes, JBs had the largest body size (Fig. 1b, Table 3), followed by the CRFs, FRFs and FEJs, respectively, when reared at L-LD (the populations showed ∼21.63%, ∼34.79% and ∼69.02% reduction in body size relative to the JBs, respectively). JBs also had the highest fecundity, followed by CRFs, FRFs, and FEJs, respectively. Post-hoc tests indicated that only the fecundity of CRFs and FRFs did not significantly differ (Fig. 1a; Table 1). JBs, CRFs and FRFs showed substantial reductions in body size when subjected to larval crowding (∼66.1%, ∼70% and ∼62.93% decrease, respectively) relative to their counterparts reared at L-LD. (Fig. S1). The FEJs were an exception, wherein both body size and fecundity were not significantly altered upon experiencing larval crowding (Figs. 3 & S1).

**Figure 1:**
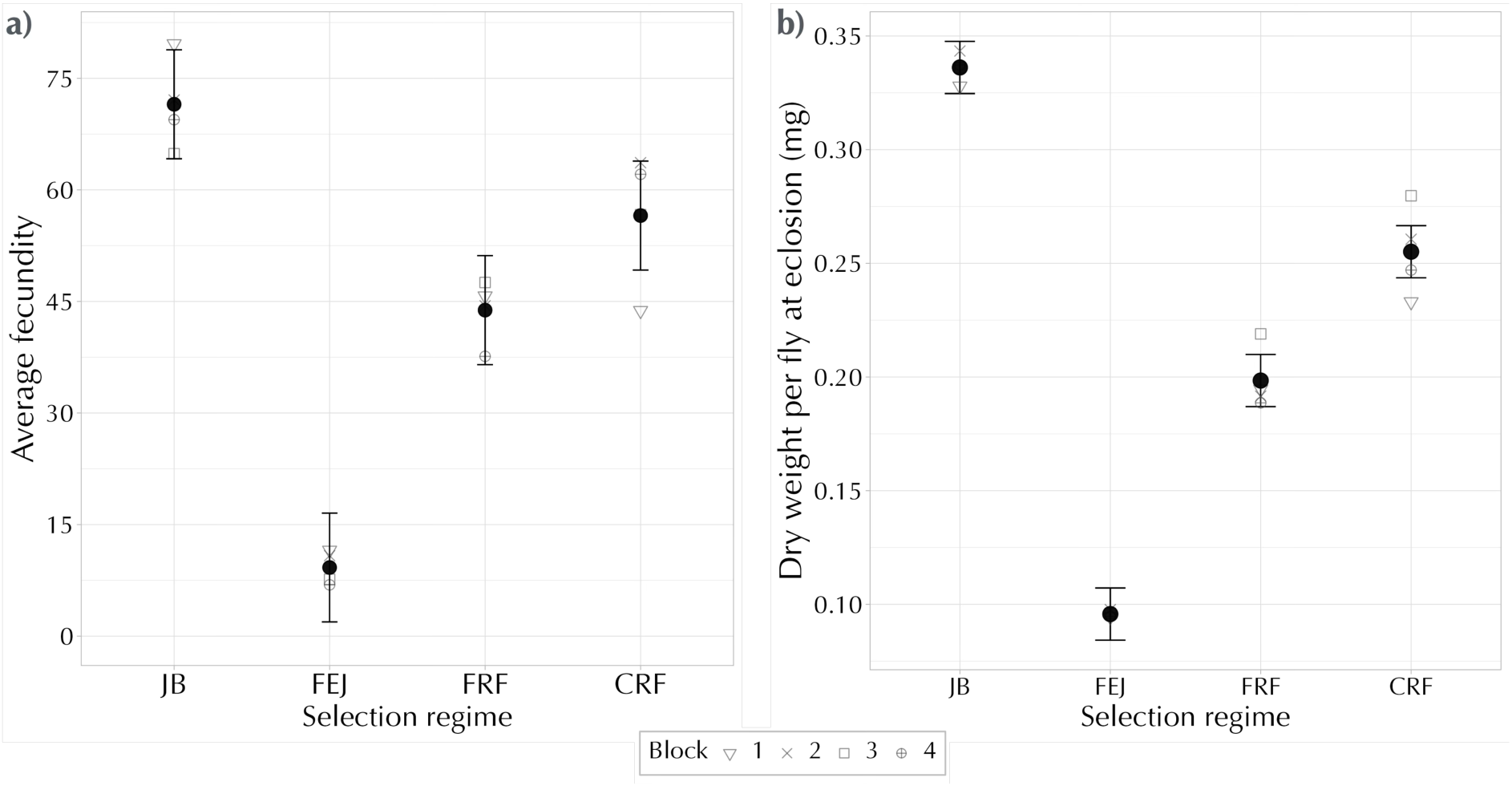
**a)** Mean fecundity of female flies of each selection regime after experiencing adult conditioning (averaged over other factors) **b)** Mean dry weight at eclosion for flies reared at low larval density (averaged over sex). The error bars represent 95% confidence intervals around the means and can be used for visual hypothesis testing.

### Smaller flies experience lower mortality costs when subjected to adult crowding

In our study, we considered mortality to be indicative of the stress experienced during the adult conditioning period, which could also subsequently affect the fecundity of females.

Mortality in females was significantly higher than in males across all selection regimes (Table 2, main effect of sex, *F*_1, 3_ = 62.76, *p* = 0.004), likely due to their larger body size and increased activity at the food surface relative to the males. Interestingly, we found that the mortality incurred depended on the size of the flies. On average, flies reared at L-LD (large flies) experienced significantly higher mortality (∼13.3% higher, mostly female) at high adult density (H-AD) compared to low adult density (L-AD). In contrast, flies reared at H-LD (small flies) showed minimal difference in mortality (∼3%) across the two adult conditioning densities (Fig. 2; Table 2, the interaction between larval and adult density, *F*_1, 3_ = 54.15, *p* = 0.005).

**Figure 2:**
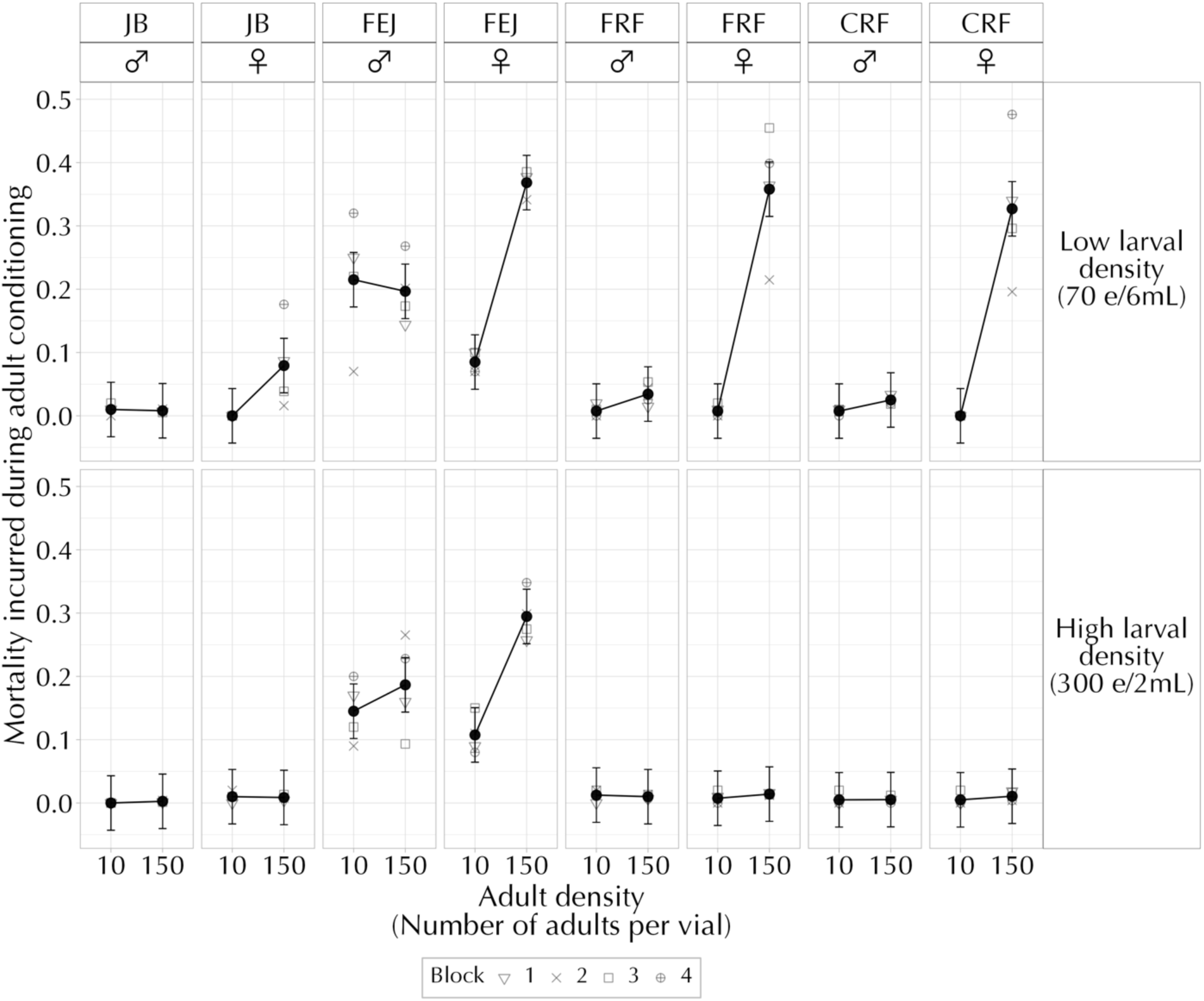
Mean proportion of mortality incurred during the adult conditioning for all combinations of selection regime, sex, larval and adult density (averaged over day of measurement). The error bars represent 95% confidence intervals around the means and can be used for visual hypothesis testing.

**Table 2.**
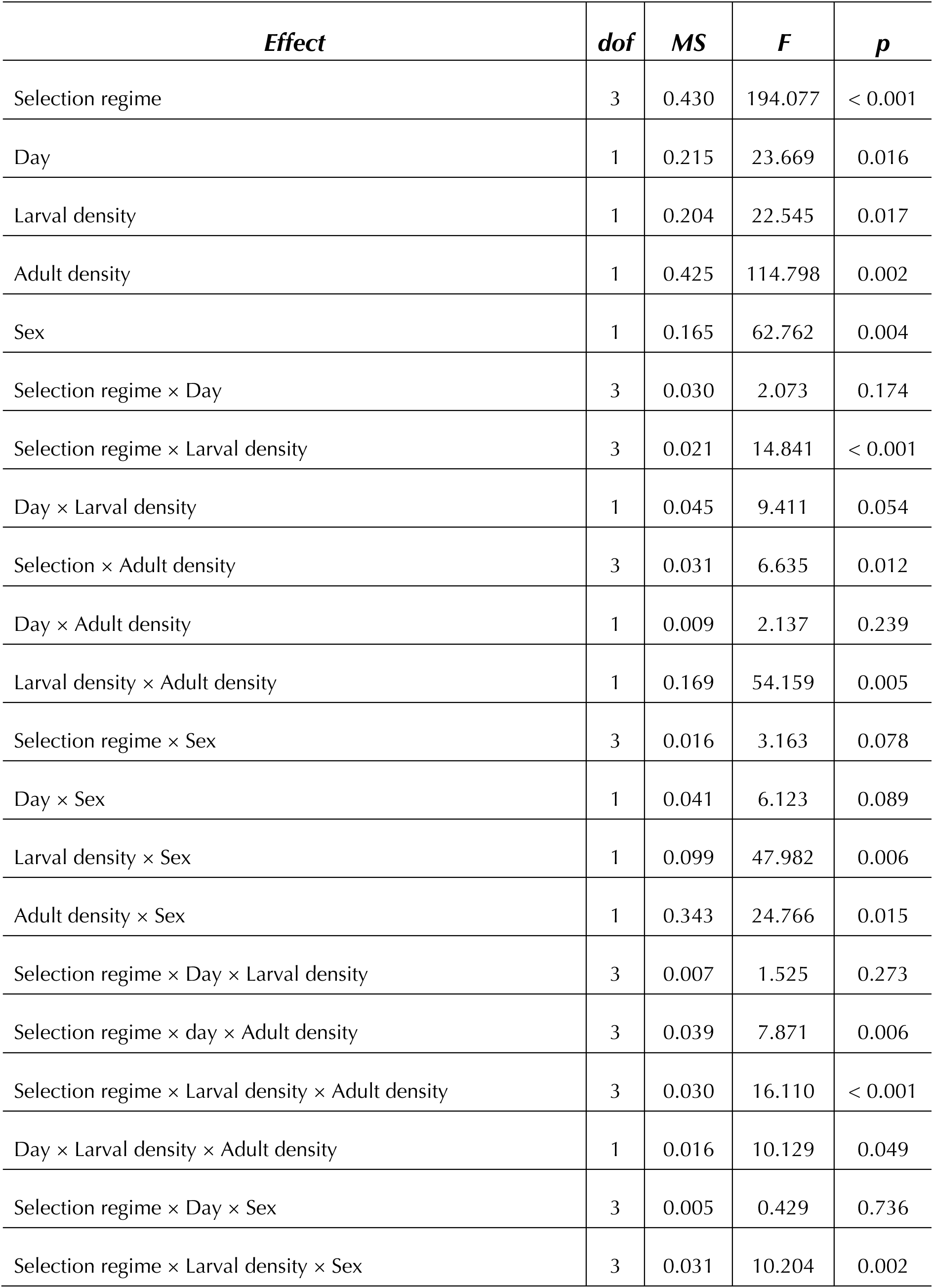
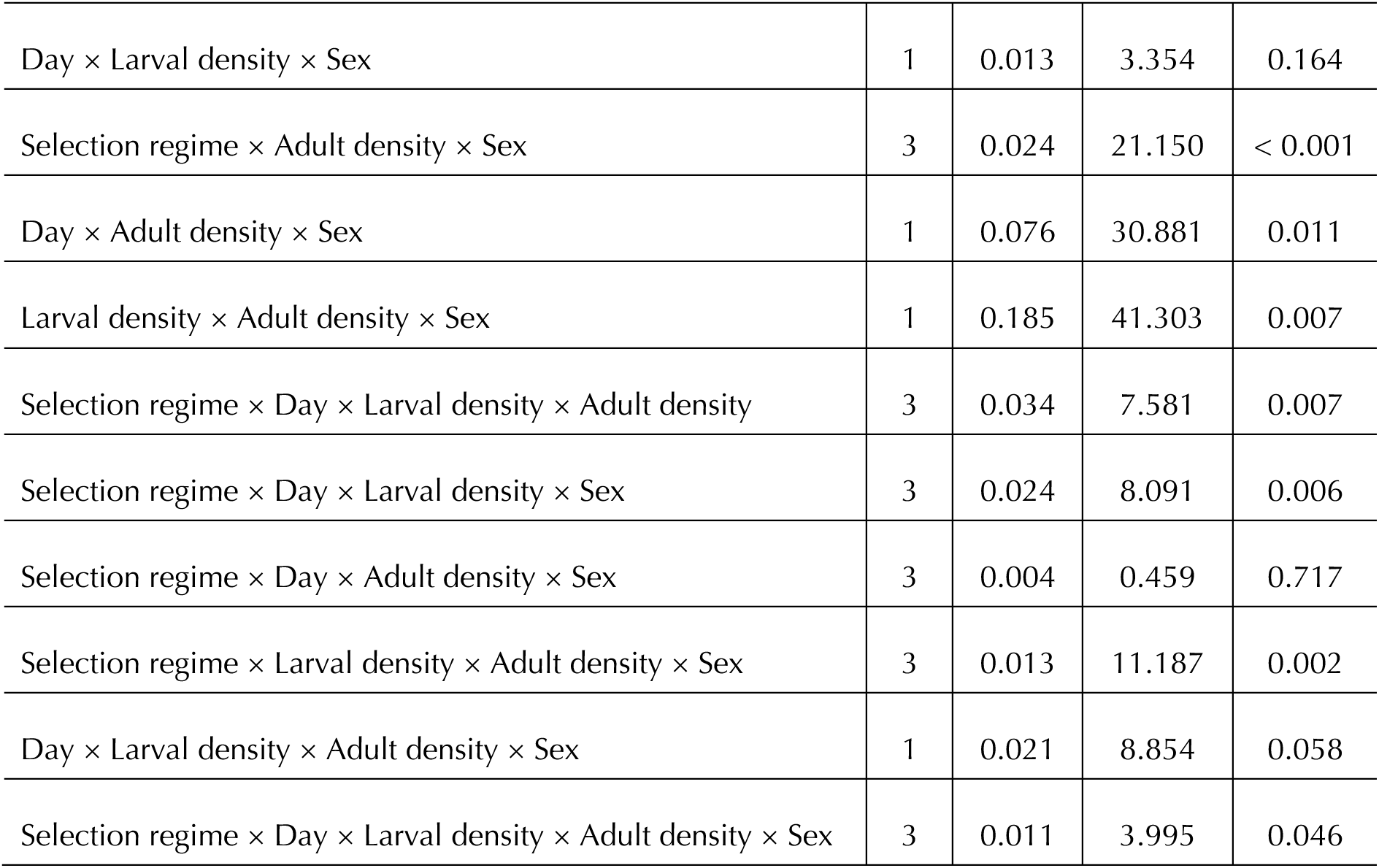
ANOVA results for mortality incurred during the adult conditioning period. In this design, random factors and their interactions are not tested for significance and are omitted from the table.

| *Effect* | *dof* | *MS* | *F* | *p* |
| --- | --- | --- | --- | --- |
| Selection regime | 3 | 0.430 | 194.077 | < 0.001 |
| Day | 1 | 0.215 | 23.669 | 0.016 |
| Larval density | 1 | 0.204 | 22.545 | 0.017 |
| Adult density | 1 | 0.425 | 114.798 | 0.002 |
| Sex | 1 | 0.165 | 62.762 | 0.004 |
| Selection regime × Day | 3 | 0.030 | 2.073 | 0.174 |
| Selection regime × Larval density | 3 | 0.021 | 14.841 | < 0.001 |
| Day × Larval density | 1 | 0.045 | 9.411 | 0.054 |
| Selection × Adult density | 3 | 0.031 | 6.635 | 0.012 |
| Day × Adult density | 1 | 0.009 | 2.137 | 0.239 |
| Larval density × Adult density | 1 | 0.169 | 54.159 | 0.005 |
| Selection regime × Sex | 3 | 0.016 | 3.163 | 0.078 |
| Day × Sex | 1 | 0.041 | 6.123 | 0.089 |
| Larval density × Sex | 1 | 0.099 | 47.982 | 0.006 |
| Adult density × Sex | 1 | 0.343 | 24.766 | 0.015 |
| Selection regime × Day × Larval density | 3 | 0.007 | 1.525 | 0.273 |
| Selection regime × day × Adult density | 3 | 0.039 | 7.871 | 0.006 |
| Selection regime × Larval density × Adult density | 3 | 0.030 | 16.110 | < 0.001 |
| Day × Larval density × Adult density | 1 | 0.016 | 10.129 | 0.049 |
| Selection regime × Day × Sex | 3 | 0.005 | 0.429 | 0.736 |
| Selection regime × Larval density × Sex | 3 | 0.031 | 10.204 | 0.002 |
| Day × Larval density × Sex | 1 | 0.013 | 3.354 | 0.164 |
| Selection regime × Adult density × Sex | 3 | 0.024 | 21.150 | < 0.001 |
| Day × Adult density × Sex | 1 | 0.076 | 30.881 | 0.011 |
| Larval density × Adult density × Sex | 1 | 0.185 | 41.303 | 0.007 |
| Selection regime × Day × Larval density × Adult density | 3 | 0.034 | 7.581 | 0.007 |
| Selection regime × Day × Larval density × Sex | 3 | 0.024 | 8.091 | 0.006 |
| Selection regime × Day × Adult density × Sex | 3 | 0.004 | 0.459 | 0.717 |
| Selection regime × Larval density × Adult density × Sex | 3 | 0.013 | 11.187 | 0.002 |
| Day × Larval density × Adult density × Sex | 1 | 0.021 | 8.854 | 0.058 |
| Selection regime × Day × Larval density × Adult density × Sex | 3 | 0.011 | 3.995 | 0.046 |

**Table 3.** ANOVA results for dry weight at eclosion for flies reared at low-larval density (70 eggs in 6mL of food). In this design, random factors and their interactions are not tested for significance and are omitted from the table.

| *Effect* | *dof* | *Msq* | *F* | *p* |
| --- | --- | --- | --- | --- |
| Selection | 3 | 0.081 | 377.101 | < 0.001 |
| Sex | 1 | 0.023 | 523.159 | < 0.001 |
| Selection × Sex | 3 | 0.001 | 77.695 | < 0.001 |

Large flies (reared at L-LD) of all selection regimes, except JBs, suffered significantly higher mortality at H-AD relative to L-AD (Fig. 2; Table 2, the interaction effect of selection regime, larval and adult density, *F*_3, 9_ = 16.11, *p* < 0.001). Interestingly, despite being the largest, JBs incurred very low mortality across both adult conditioning densities, likely due to inadvertent selection for adaptations to mild adult crowding (see Discussion). In comparison, the FEJs showed consistently high mortality across all treatment combinations despite being the smallest in body size. The elevated mortality of FEJs is likely a result of the selection pressures they experience (see Discussion).

### Small flies show an increase in female fecundity in response to an increase in adult density

Fecundity varied based on the combination of larval and adult density experienced by the flies. For large flies (reared at L-LD), we observed significantly lower fecundity (∼17.3% lower) when exposed to H-AD, indicating a detrimental effect of experiencing adult crowding (Fig. 3, Table 1, the interaction between larval and adult density, *F*_1, 3_ = 2966, *p* < 0.001). In contrast, small flies (reared at H-LD) showed higher fecundity (∼24.1% higher) with an increase in adult density (Fig. 3).

**Figure 3:**
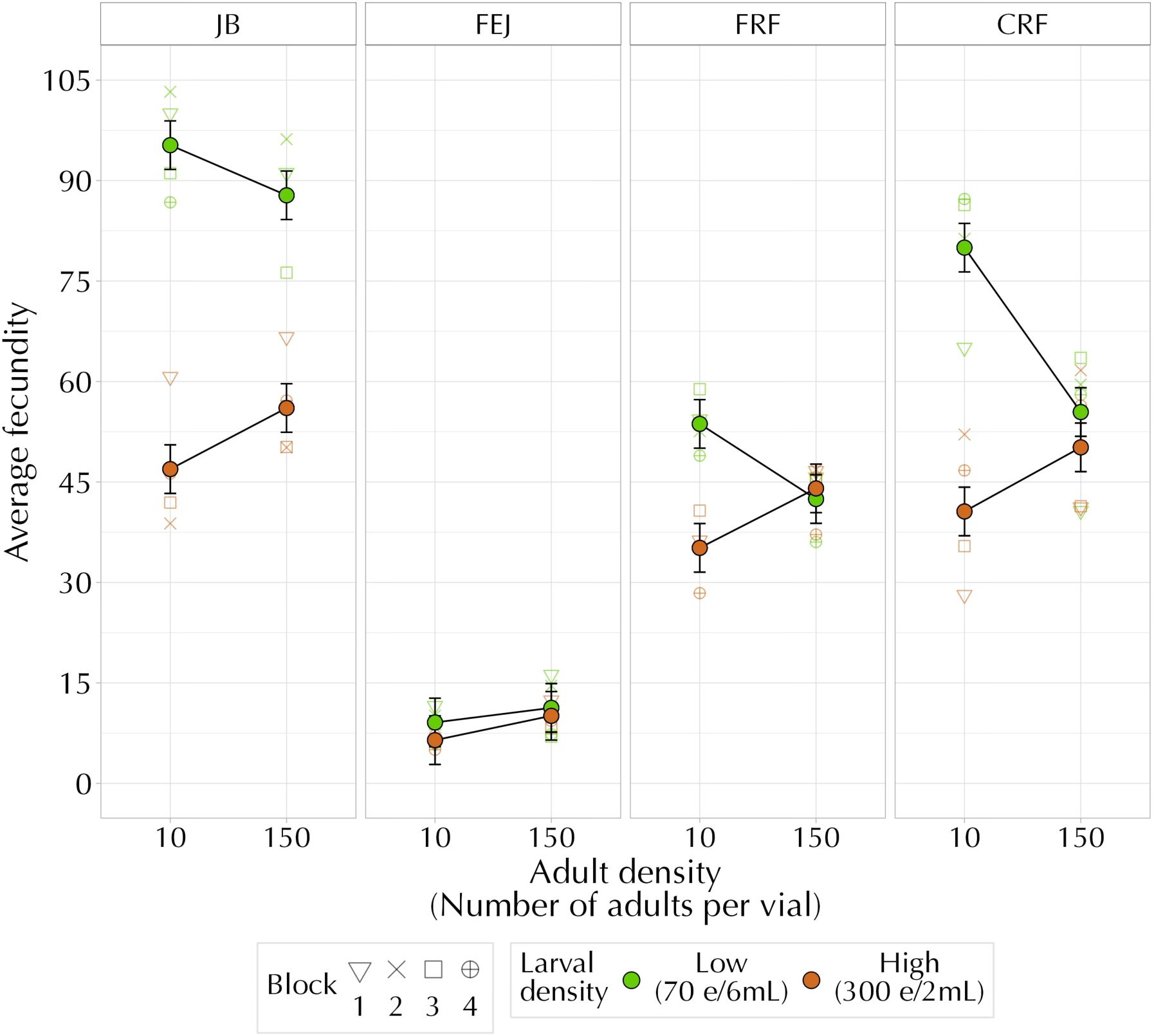
Mean female fecundity after experiencing adult conditioning for all combinations of selection regime, larval and adult density (averaged over day of measurement). The error bars represent 95% confidence intervals around the means and can be used for visual hypothesis testing.

Interestingly, the FEJs showed an increase in fecundity with an increase in adult density regardless of the larval densities they were reared at, presumably because their resultant body size was small across both larval density treatments. In contrast, JBs, CRFs and FRFs showed a similar response of increased fecundity only when reared at H-LD (Fig. 3). When flies from these selection regimes were reared at H-LD, the resulting body sizes were similar to those of the FEJs (Fig. S1). The pattern of an increase in fecundity with an increase in adult density for the small flies remained remarkably consistent across the four replicate blocks, highlighting the robustness of the results (Fig. S2).

Our results thus suggest that smaller flies, achieved either through larval crowding (ecological effects) or as a consequence of selection pressures (as in the FEJs, evolutionary effects), experience increased fecundity after an episode of high adult density.

## Discussion

Adult crowding has typically been viewed as being detrimental to key life-history traits such as mortality, fertility and female fecundity. Our study challenges this existing view by showing that body size can mediate these effects of adult crowding in different directions (Fig. 4). We subjected flies of various body sizes to a short duration (two days) of adult crowding and observed nuanced responses that depended significantly on body size. Small flies had a consistent advantage in H-AD conditions, exhibiting lower mortality and increased fecundity than large flies when subjected to the same adult density. Our findings emphasize how body size can influence outcomes on fitness components (mortality and fecundity) when exposed to the stress of adult crowding. To our knowledge, this is the first study to directly test the interaction between adult density and body size on fitness components, with body size being manipulated by both ecological (larval density) and evolutionary (selection) means.

**Figure 4.**
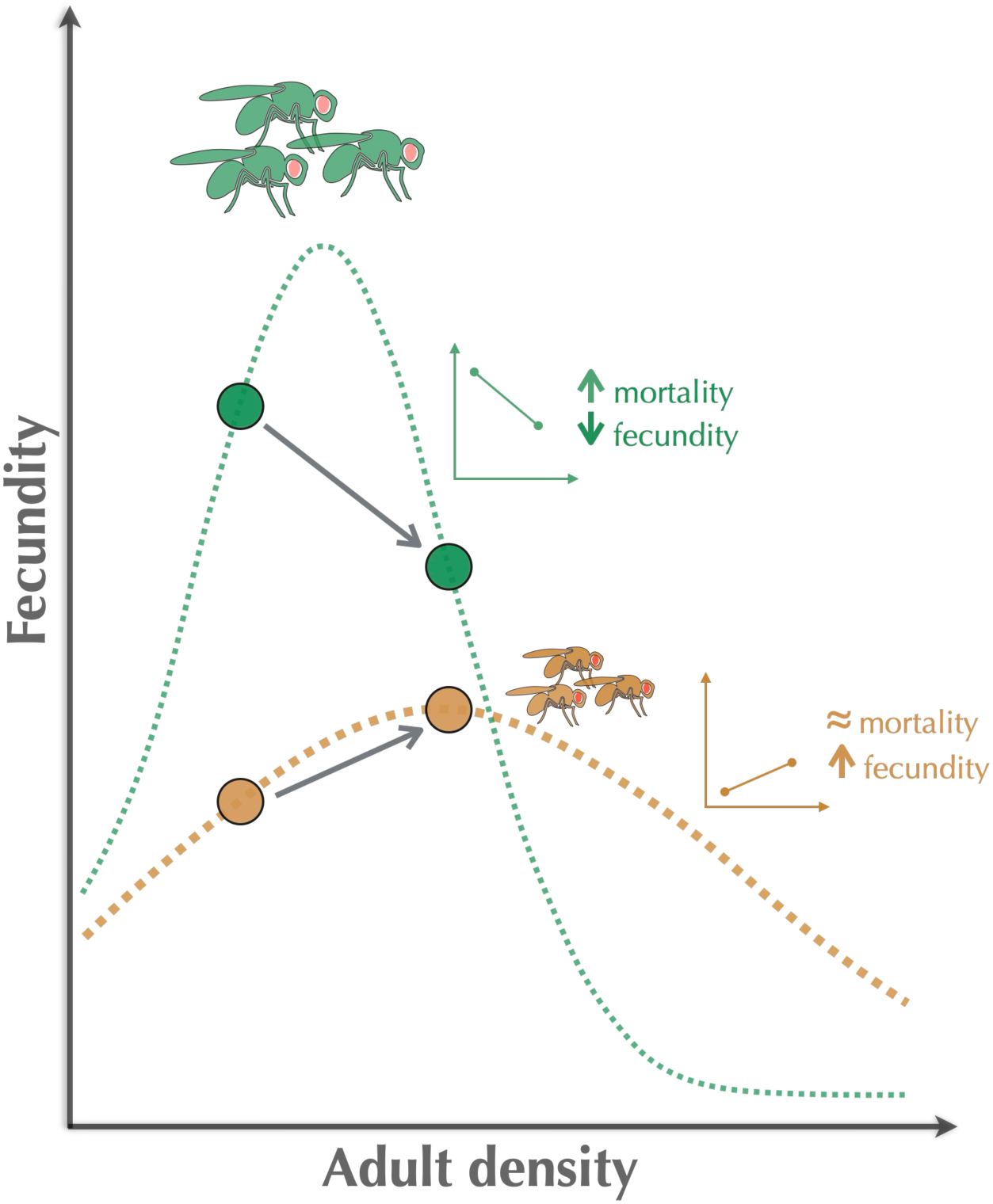
A proposed relationship between female fecundity and adult density. The model illustrates the potential relationship between female fecundity and adult density, highlighting the differences between small and large flies. Large flies show a narrow fecundity distribution with a sharp peak. In comparison, the small flies have a broader distribution, with a greater standard deviation around the mean and a flatter peak. The points highlighted on the curves represent the two adult conditioning densities used in our experiment, with corresponding patterns observed for fecundity and mortality shown alongside for comparison. The model presented here represents one possible scenario. Further studies are required to test this hypothesis as a potential explanation for the observed results.

### The role of body size

Our findings suggest that a smaller body size increases resilience under adult crowding. We observed lower mortality in small flies under H-AD than in large flies (Fig. 2), reinforcing our hypothesis that a smaller body size mitigates some physiological stress of crowding. Smaller flies occupy less space, likely reducing harmful physical interactions among the flies and creating a less stressful environment within the vial during crowding. Therefore, the adverse effects experienced would likely be fewer, leading to low mortality.

Flies from the FEJ regime consistently respond with an increase in fecundity as adult density increases, regardless of larval rearing density (Fig. 3). Other selection regimes (FRF, CRF and JB) match this response only when reared under H-LD conditions. At H-LD, the body sizes of the flies of these selection regimes are very similar to FEJs (Fig. S1), possibly eliciting the same response to adult crowding.

Notably, the advantages of small body size are evident regardless of how the size was achieved— whether through ecological effects (via larval crowding) or as an evolutionary consequence of selection pressures (as in the case of the FEJs). The FEJs, which are intrinsically small due to selection for rapid development, consistently had increased fecundity under H-AD conditions. It suggests that body size may be a key determinant of the response to adult crowding, and the pattern of increase in fecundity is not merely an artifact of larval crowding.

### Potential mechanisms driving the observed patterns

#### Negative effects of adult crowding

1. **Nutritional interference by the flies:** Under crowded conditions, adult flies can face challenges accessing the surface of the food. It can lead to individuals acquiring less nutrition during the adult crowding period. In particular, for females, insufficient nutritional status is known to affect fecundity negatively (Robertson & Sang, 1944a,b).
2. **Space constraints:** For large flies, space limitations during crowding may increase the chances of collision between flies and cause physiological stress, leading to the high mortality observed at H-AD. Additionally, females would face challenges in accessing oviposition sites at the food surface. Repeated visits to the food surface to obtain nutrition or to gain access to oviposition sites can make females more prone to physical harm due to trampling/collisions in crowded conditions (Joshi, 1997) and may even result in higher female mortality than males (Fig. 2). Higher levels of male harassment during the conditioning period could also lead to an increase in female mortality (Fowler & Partridge, 1989; Kuijper *et al*., 2006).

#### Positive effects of adult crowding

1. **Increase in matings:** Increased frequency of encounters between males and females at H-AD could facilitate an increase in matings. In the short term, multiple matings have been shown to boost the female fecundity of *Drosophila* (Hanson & Ferris, 1929; Turner & Anderson, 1983; Chen *et al*., 1988; Taylor *et al*., 2008). Additionally, there is evidence that yeasted females tend to remate more often than un-yeasted females (Trevitt *et al*., 1988). Flies in our assay were provided with yeast during the adult conditioning period, which could have contributed to increased matings.
2. **Changes in male behaviour:** Exposure to H-AD can trigger behavioural changes in males, primarily driven by the perception of increased competition. These changes can include an increase in the mating duration or an increase in the seminal fluid proteins (SFPs) transferred to the females. Consequently, these SFPs are known to temporarily boost the fecundity of females (Bretman *et al*., 2009; Wigby *et al*., 2009; Bretman *et al*., 2010; Fedorka *et al*., 2011; Bretman *et al*., 2012; Hopkins *et al*., 2019).

The interplay of these effects could shape the net effects of adult crowding on mortality and female fecundity. We considered adult mortality that occurred during the conditioning period to reflect the degree of stress experienced by the flies, with higher mortality signifying higher levels of stress experienced. Overall, adult crowding positively affected the small flies (no change in mortality and an increase in fecundity compared to L-AD conditions).

### Impact of selection pressures

Given the differences in body size among the four selection regimes (Fig. 1b), we initially hypothesized that larger individuals would exhibit higher mortality with increased adult density, with the mortality pattern of JBs > CRFs > FRFs > FEJs in L-LD and H-AD treatment combinations. However, the observed mortality pattern was FEJs > FRFs ≈ CRFs > JBs. Despite having the smallest body size, the FEJs had the highest mortality at H-AD conditions, regardless of the larval-rearing density. The mortality levels were similar for males and females (Fig. 2), unlike the typical pattern of higher female mortality observed across other selection regimes.

The elevated mortality levels of FEJs are not surprising, and we attribute this to their generally poor performance under various stressful conditions. Previous studies have shown that FEJs have low egg-to-adult survivorship under stressful conditions created by larval crowding, starvation and increased waste content in their food (Prasad, 2003). Moreover, the FEJs appear to have evolved poor levels of physical activity (personal observations), likely due to very low energy reserves as adults (Prasad, 2003). For instance, when reared at L-LD, several freshly eclosed FEJ adults often stick to the food surface and die, a phenomenon rarely observed in the other selection regimes. This mortality is not due to crowding effects within the vial per se but because FEJ flies find it hard to extricate themselves from soggy food, if stuck. The CRFs and FRFs, derived from the FEJs via the relaxation of these selection pressures, have been under these relaxed conditions for over 100 generations. Certain aspects of the inherent poor levels of physical activity of the FEJs may persist in the CRFs and FRFs and may not have returned to the ancestral state of JBs. These evolved constraints may play out particularly under stressful conditions and contribute to higher mortality at H-AD relative to the JBs.

Being the largest in size (Fig. 1b), JBs were expected to incur high levels of mortality after being subjected to H-AD. However, we observed low mortality in combinations of L-LD and H-AD compared to other selection regimes (Fig. 2). This can be due to inadvertent selection for mild adult crowding imposed on the JBs as part of routine maintenance. During laboratory maintenance, JBs undergo three rounds of vial-to-vial transfers after adults eclose. After one round of vial-to-vial transfers, flies are housed in the vials for 2 days before receiving fresh food through another round of vial-to-vial transfers. The density within each vial is ∼50-70 adults. The ancestors of JBs, the UUs (first described in Mueller *et al*., 1993), which were in turn derived from the Bs (Rose, 1984), and their ancestors, the Ives populations (Rose & Charlesworth, 1981), were also maintained with vial-to-vial transfers. Therefore, the population has been maintained in vials for over 1000 generations. This period within the vials could be considered as conditioning with mild levels of adult crowding, and over generations, it may have conferred resilience to the adult crowding experienced in our assay. These findings underscore the importance of considering the evolutionary history and the prevalence of any inadvertent selection pressures when interpreting results from evolutionary experiments.

For the FEJs, the smaller body size has been achieved via the evolution of reduced minimum critical size relative to the JBs (Prasad *et al*., 2001), and the larvae leave the food soon after attaining this size. In JBs, larvae can feed long after attaining the minimum critical size, making the adults significantly larger than the FEJs. Consequently, unlike other selection regimes, the FEJs do not show a large decrease in body size under larval crowding (Fig. S1), underscoring the evolutionary constraints operating on their body size. In *Drosophila*, female fecundity positively correlates to body size (Robertson, 1957; Lefranc & Bundgaard, 2000; Byrne & Rice, 2006). The constraints on body size in FEJs could, in turn, impose a constraint on female fecundity and explain why they do not show a significant increase in fecundity when exposed to H-AD treatments. It is also known that FEJ females allocate relatively less lipid to early-life reproduction than JB females (Prasad, 2003).

### Broader implications of our study

Our findings challenge the long-standing assumption that adult crowding is universally detrimental, instead revealing that its effects are mediated by body size. Earlier studies on adult crowding typically used flies reared under L-LD conditions, and so, as the adults were large, we speculate that the considerable adverse effects of crowding reduced the fecundity of the surviving females. The consistent use of large flies (reared at L-LD) across earlier experiments that examined the effects of adult crowding led to the view that adult crowding typically induced high mortality and reduced the fecundity of surviving females. In this study, a significant interaction effect was observed between larval density (which impacts the adult body size) and adult density for both mortality and fecundity (Tables 1 & 2). We highlighted how smaller body sizes can shift the balance of effects experienced during adult crowding toward positive outcomes. Small flies consistently showed increased fecundity under H-AD (relative to L-AD), while large flies (reared under L-LD) had lowered fecundity at H-AD relative to L-AD. As previous studies typically used flies reared at L-LD, their response to adult crowding remained largely negative.

We hypothesize that there is a hump-shaped relationship between adult density and fecundity. We propose that the response norm will be altered based on the body size of the flies subjected to varying adult densities (Fig. 4). For large flies, fecundity initially increases with adult density but declines beyond a certain threshold of density. For small flies, the threshold would lie at a much higher adult density, so the increase in fecundity would be observed over a broader range of adult densities. In our experiment, the two adult conditioning density treatments perhaps captured the response falling on either side of the curve for the large flies, resulting in lowered fecundity at H-AD. However, the small flies remained on the increasing side of the curve at the high adult conditioning density, explaining the fecundity boost at H-AD observed in our study.

### Future directions

Our results indicate that the benefits of a small body size subjected to adult crowding are not limited to incurring low mortality but also include gains in fecundity, likely driven by high mating rates. Future studies could explore the relationship between mating frequency and fecundity in crowded environments to elucidate this phenomenon further. Large flies (reared at L-LD) experienced high mortality in the H-AD treatment. On the contrary, JBs, CRFs, and FRFs reared at H-LD were significantly smaller and showed similar levels of mortality across both adult conditioning densities. These results suggest that stress levels experienced during crowding can vary depending on the body size of individuals. Small flies would occupy less space within a vial and would have more effective space available compared to the same number of large flies. Thus, the physical space within the vial, including the volume of the air column available to adults, could also influence the outcome of crowding. Altering vial dimensions and air volume (thereby, the space available for flies) could provide deeper insights into the interaction between body size and fitness under conditions of adult crowding.

Our experimental design involved the presence of a male and female during the oviposition window. A possible consequence is the negative effect a male can have on female fecundity. Sexual conflict, mainly through mate harm, is known to impact female fecundity negatively. The JB and CRF populations are known to have males that cause more harm to females upon mating relative to the other selection regimes (Mital, 2019; Mital *et al*., 2021, 2022; C. Temura & A. Joshi, unpublished data). In such cases, the levels of mate harassment experienced during the oviposition window would depend on the intensity of sexual conflict prevailing in the selection regime. Additionally, females could experience compounded fecundity costs due to mate harassment. For example, the FEJ females could be affected to a far lower degree by the presence of FEJ males, as opposed to the impact that JB males have on the female counterparts during the oviposition window, as sexual conflict is much lower in the FEJs relative to the JBs. The impact of the presence versus absence of males during oviposition on female fecundity can be further explored.

Smaller flies, whether achieved through larval crowding or as a response to selection pressures, consistently perform better under H-AD, highlighting the importance of body size in determining the response to crowding over and above the method used to obtain smaller body sizes. However, the response shown by flies reared at H-LD could be an artifact of being subjected to some form of larval density in the first place. Future studies can use a single selection regime to decouple the effects of body size from other confounding factors (involving evolutionary history or ecological methods of achieving smaller sizes).

Larger flies (the JBs) were expected to experience higher mortality levels when crowded as adults. However, we found the levels of mortality to be highest in FEJs, which were the smallest in size (Figs. 1b & 2). In this case, it would appear that the selection regimes are a confounding factor. The link between body size and fly response to adult crowding can be further elucidated by studying flies over a range of body sizes (obtained by varying larval density for a single selection regime) under adult crowded conditions.

In conclusion, our results emphasize the importance of body size in influencing patterns of mortality and fecundity when flies are subjected to a brief episode of adult crowding. Thus, the effects of adult crowding on key fitness components can be interpreted with a more nuanced perspective than the previously established view.

## Materials and Methods

### Study populations

This study used four sets of four replicate large (∼1800 breeding individuals), laboratory-maintained *D. melanogaster* populations. The Joshi Baseline (JB_1-4_) populations served as ancestral controls for the other three sets of populations. From the JBs, populations selected for faster development and early reproduction were derived (Faster developing, Early reproducing JB-derived; FEJ_1-4_). Subsequently, two sets of reverse-selected populations were derived from the FEJs. The Completely Relaxed FEJ-derived (CRF_1-4_) populations were relaxed for selection on both development time and the time of reproduction, whereas the Faster development Relaxed FEJ-derived (FRF_1-4_) populations were relaxed only for selection on development time but not for early reproduction. The maintenance protocols and ancestry for all four sets of populations have been reported in detail earlier (*For JBs:* Sheeba *et al*., 1998; *For FEJs:* Prasad *et al*., 2000; *For CRFs and FRFs*, then labelled the RFs and RRFs respectively: Mital, 2019) and are summarized below. Each subscript represents ancestral descent from a specific JB population. Hence, FEJ_1_ is ancestrally closer to JB_1_, FRF_1_ and CRF_1_ than to the other FEJ populations. Our design treats populations with the same numerical subscript as blocks (subsumes ancestry and replicate runs of the experiment) for statistical analyses. The assay was performed one block at a time, for logistic reasons.

### Maintenance protocols

All populations are maintained at 25±1°C, high humidity and constant light on banana-jaggery medium. The JBs are maintained on a 21-day discrete generation cycle, wherein eggs are collected in 8-dram glass vials (2.2 cm dia × 9.6 cm ht) at a density of 60-80 eggs in ∼6 mL of food medium. Once all adults eclose (day 12 from egg collection), they are transferred to fresh adult collection vials containing ∼4 mL of medium. Thereafter, adults undergo two more rounds of vial-to-vial transfers to receive fresh food on days 14 and 16 from egg collection. On day 18, the flies are transferred to Plexiglas cages (25 × 20 × 15 cm^3^) and provided with a Petri plate containing food medium coated with yeast paste (henceforth, yeasted food plate) for two days.

Following this, a fresh food plate cut into two halves is provided for oviposition. The vertical edges created by cutting the food stimulate oviposition in the females. Eggs from this plate are collected after 18 hours to initiate the next generation (on day 21).

For the FEJs, eggs are collected in the same manner and at the same density as the JBs. Once adults begin to eclose, only the earliest (∼20-25%) eclosing adults from each vial are transferred to Plexiglas cages, which form the breeding pool for initiating the next generation. A yeasted food plate is then provided for three days, following which, a fresh food plate with vertical edges is provided for 1-3 hours for egg-laying, which is used to initiate the next generation. Consequently, these populations have evolved a considerably shorter egg-to-adult development time and are presently maintained on a 10-day discrete generation cycle.

The CRFs and FRFs were derived from the FEJs around the 550^th^ generation of FEJ maintenance. For CRFs, the egg collection protocol is the same as for the JBs and FEJs, and all adults eclosing from the culture vials undergo vial-to-vial transfers similar to the JBs, but on days 10 and 12 from egg collection. Following this, they are transferred to Plexiglas cages on day 14 and provided with a yeasted food plate for two days. The eggs to initiate the next generation are collected on day 17 (similar to the protocol for JBs). They are presently maintained on a 17-day discrete generation cycle in the laboratory. Thus, in CRFs, selection for rapid development and early reproduction have both been relaxed.

In the case of FRFs, relaxed only for selection for rapid development, while the egg collection protocol remains the same, all eclosing adults are transferred directly to the cages on day 9 from egg collection and are then provided with a yeasted food plate for two days before initiating the next generation on day 12 (similar to the protocol for JBs). They are presently maintained on a 12-day discrete generation cycle. The CRFs and FRFs, therefore, differ in the amount of breeding time available to the adults, and any evolved differences between them can thus be ascribed to differences in the breeding time, whereas any differences found between the FRFs and FEJs can be ascribed to the role of selection for rapid development.

At the time of our assay, the JBs had been maintained in the laboratory for 385 generations, FEJs had undergone 755 generations of forward selection, and the CRFs and FRFs had undergone 131 and 195 generations of reverse selection, respectively.

### Standardization of populations

Before starting the assay, we imposed one generation of standardized/common rearing conditions across all selection regimes to eliminate non-genetic parental effects (Rose, 1984). For this, we collected eggs from all sixteen populations at a density of ∼70 eggs in 6 mL of banana-jaggery medium. As the four selection regimes differ in egg-to-adult development times, the larval cultures initiated from each selection regime were staggered to ensure that adults emerged together. All eclosed flies were transferred to a Plexiglas cage, and a yeasted food plate was provided. After two days, the flies were given a fresh food plate cut into two halves for egg laying. After 18 hours, eggs were collected from these plates for the assay egg collection. The flies used in the assay were derived from one of two larval densities: Low (70 ± 10 eggs in 6 mL of food) and high (300 ± 20 eggs in 2 mL of food).

### Collection of eclosing flies

Flies from the low larval density (L-LD) treatments were transferred to Plexiglas cages once all adults eclosed (between days seven and eleven from assay egg collection, depending on the selection regime), For the high larval density vials (H-LD), the eclosion of adults was spread out over several days from egg collection (between days six and fifteen). These flies were transferred once daily to a Plexiglas cage for every population. Separate cages were maintained for flies whose fecundity was measured on the 12^th^ and 18^th^ day from egg collection and freshly eclosed flies were transferred to cages till the day of the adult conditioning setup (10^th^ and 16^th^ day, respectively). In all cages, flies were provided with fresh food plates on alternate days from the start of eclosion till the setup of adult conditioning vials.

### Adult crowding

Flies were exposed to two kinds of adult conditioning densities for two days *before* their fecundity was measured: Low adult-density (L-AD) vials had five males and five females per vial, and high adult-density (H-AD) vials had 75 males and 75 females per vial. All vials were provided with ad-libitum yeast paste applied onto one side of the wall, along with ∼4 mL of banana-jaggery medium. Flies from the holding cages were introduced into the vials using aspirators to avoid exposing the flies to CO_2_ before the adult conditioning period. Ten and five replicate vials were set up for the low and high adult density conditioning treatments respectively for each treatment combination of selection regime, larval density, day of measurement and replicate population.

To facilitate a comparison of effects that also correlated reasonably well with regular maintenance protocols of the populations, we measured fecundity at two different time points - 12 days from egg-lay (optimal in terms of the FEJ and FRF maintenance protocols) and 18 days from egg-lay (optimal in terms of the CRF and JB maintenance protocols). Thus, for flies whose fecundity was measured on day 12 from egg collection, the individuals experienced the adult conditioning densities on days 10 and 11 from egg collection. Similarly, for flies whose fecundity was measured on day 18, the individuals experienced the adult conditioning treatment on days 16 and 17 from egg collection.

### Mortality and fecundity measurements

Sex-specific mortality for each vial was recorded at the end of the conditioning period. Surviving males and females from each treatment combination were used to measure fecundity. Using an aspirator, twenty male-female pairs per treatment combination were set up as one male-female pair per vial with ∼0.5 mL of food medium (just enough to cover the base of the vial, to facilitate accurate egg counts). After 24 hours, flies were discarded and the number of eggs in each vial was counted.

### Dry weight measurements

The dry weight measurements were obtained from an earlier, independent assay (M. Rao, & A. Joshi, unpublished data). All sixteen populations were subjected to one generation of standardization prior to the assay, as detailed above. The adults of the standardized populations were provided with food plates cut into two halves for egg-laying. These plates were prepared with twice the usual amount of agar to facilitate easy separation of eggs, in order to count the precise number of eggs required for the assay. After 1 hour, eggs were collected from the plates at a density of exactly 70 eggs in 6 mL of food, with ten replicate vials for each selection regime × block combination. Freshly eclosed flies from each culture vial were transferred into empty vials, frozen and stored at −20℃. The flies were separated by sex and dried in a hot-air oven at 70℃ for 36 hours. Following this, they were divided into five batches of five flies each for all treatment combinations. Each batch of flies underwent triplicate measurements on a Sartorius (CP 225D) fine balance, and the average of these readings was taken as the weight of the batch. At the time of the assay, JBs had undergone 371 generations of laboratory maintenance, FEJs had undergone 728 generations of forward selection, and the CRFs and FRFs had undergone 115 and 173 generations of reverse selection, respectively.

### Statistical analyses

We carried out a full-factorial, mixed model Analyses of Variance (ANOVAs) on the adult mortality, female fecundity and dry weight data obtained from the experiments. We treated the selection regime, day of measurement, larval density, and adult conditioning density as fixed factors in the analysis for female fecundity. For the adult mortality ANOVA, selection regime, day of measurement, sex, larval density and adult density were considered as fixed factors. For the dry weight ANOVA, selection regime and sex were considered as fixed factors. In all the analyses, the replicate populations were treated as a random block factor, crossed with the other factors. The cell means for each treatment combination were used as the units of analysis. Pairwise post-hoc comparisons were carried out using Tukey’s HSD.

All ANOVAs were run on R release 4.4.1 using the *stats* package (by calling upon the *aov* function) (R Core Team 2025). All graphs were plotted using R release 4.4.1 using the packages *tidyverse* and *ggplot2* (Wickham, 2016; Wickham *et al*., 2019).

## Acknowledgements

We thank Srikant Venkitachalam, Bhavya Pratap Singh, Sajith V.S., Ramesh K., Rajanna N. and Muniraju P. for their assistance in the laboratory. MR was supported by a doctoral fellowship from JNCASR. The study was supported by intramural funds from JNCASR, a Science and Engineering Research Board (SERB) JC Bose National Fellowship (recipient: AJ), and, in part, by A. Joshi’s personal funds.

## Author contributions

MR, AM and AJ conceptualized this study and designed the experiment; MR, AM, CT and AS carried out the experiments; MR analyzed the data and wrote the first draft of the paper; MR, CT and AJ contributed to subsequent revisions and the final draft of the paper.

## Notes

### Competing Interest Statement

The authors have declared no competing interest.

