## Supplementary Materials for "Bigger is not always better: size-dependent fitness effects of adult crowding in *Drosophila melanogaster*"

#### Supplementary Text

##### Text S1: Experiment to obtain dry weights of flies shown in Figure S1

The dry weight measurements were obtained from an independent pilot assay. All selection regimes were subjected to one generation of standardization, as described earlier. The adults of the standardized populations were provided with yeasted food plates for two days, and then a food plate cut into two halves for egg-laying. After 18 hours, eggs were collected from the plates at two types of larval densities per selection regime – a low larval density (70 ± 10 eggs in 6 mL of food) and high larval density (300 ± 20 eggs in 2 mL of food). As adults eclosed from the culture vials, they were transferred daily into empty vials, frozen and stored at -20°C. Flies were separated by sex and dried in a hot-air oven at 70°C for 36 hours. Following this, they were divided into five batches of ten flies each. Each batch of flies underwent triplicate measurements on a Sartorius (CP 225D) fine balance, and the average of these readings was taken as the weight of the batch. At the time of the assay, JBs had undergone 459 generations of laboratory maintenance, FEJs had undergone 908 generations of forward selection, and the CRFs and FRFs had undergone 222 and 322 generations of reverse selection, respectively. As dry weights from only two replicate blocks were measured, the data is to be interpreted with caution.

### Supplementary Tables

**Table S1 -** ANOVA results for dry body weight of the flies at eclosion. In this design, random factors and their interactions are not tested for significance and are omitted from the table.

| ***Effect*** | ***dof*** | ***Msq*** | ***F*** | ***p*** |
|---|---|---|---|---|
| Selection | 3 | 2.191 | 32.131 | 0.008 |
| Sex | 1 | 1.043 | 2344.39 | 0.013 |
| Larval density | 1 | 14.561 | 226.172 | 0.042 |
| Selection × Sex | 3 | 0.042 | 3.729 | 0.154 |
| Selection × Larval density | 3 | 1.313 | 46.463 | 0.005 |
| Sex × Larval density | 1 | 0.576 | 369.672 | 0.033 |
| Selection × Sex × Larval density | 3 | 0.037 | 47.439 | 0.005 |

### Supplementary Figures

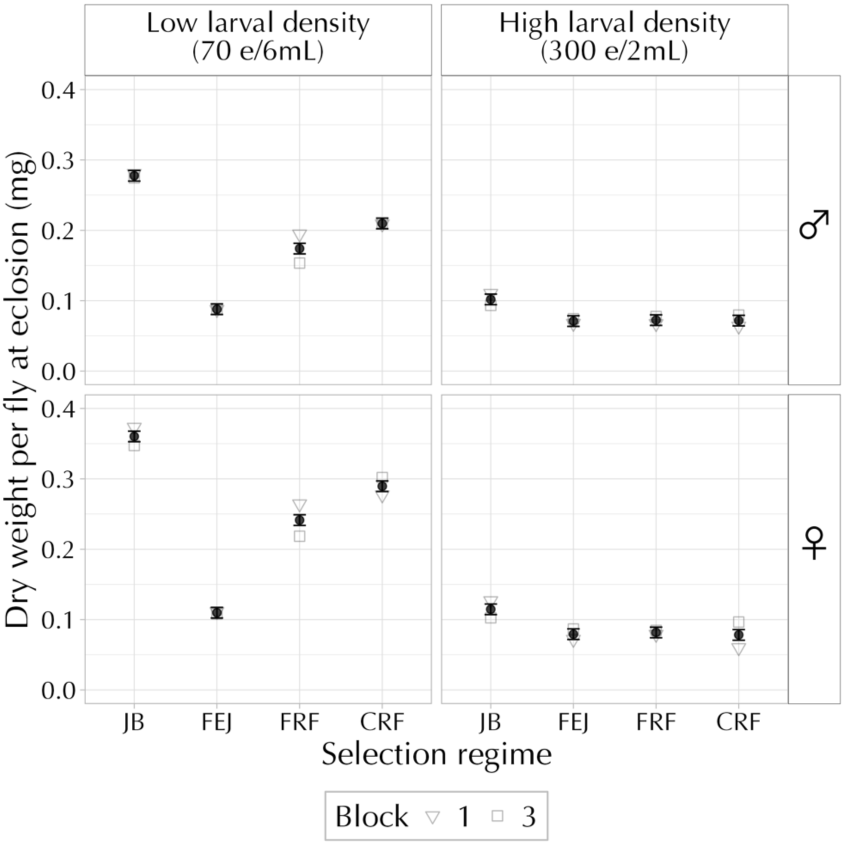


**Figure S1:** Mean dry weight at eclosion for flies from all selection regimes reared at two larval densities. The data for this figure involves only two replicate blocks, and therefore, results must be interpreted with caution. The error bars represent 95% confidence intervals around the means and can be used for visual hypothesis testing.

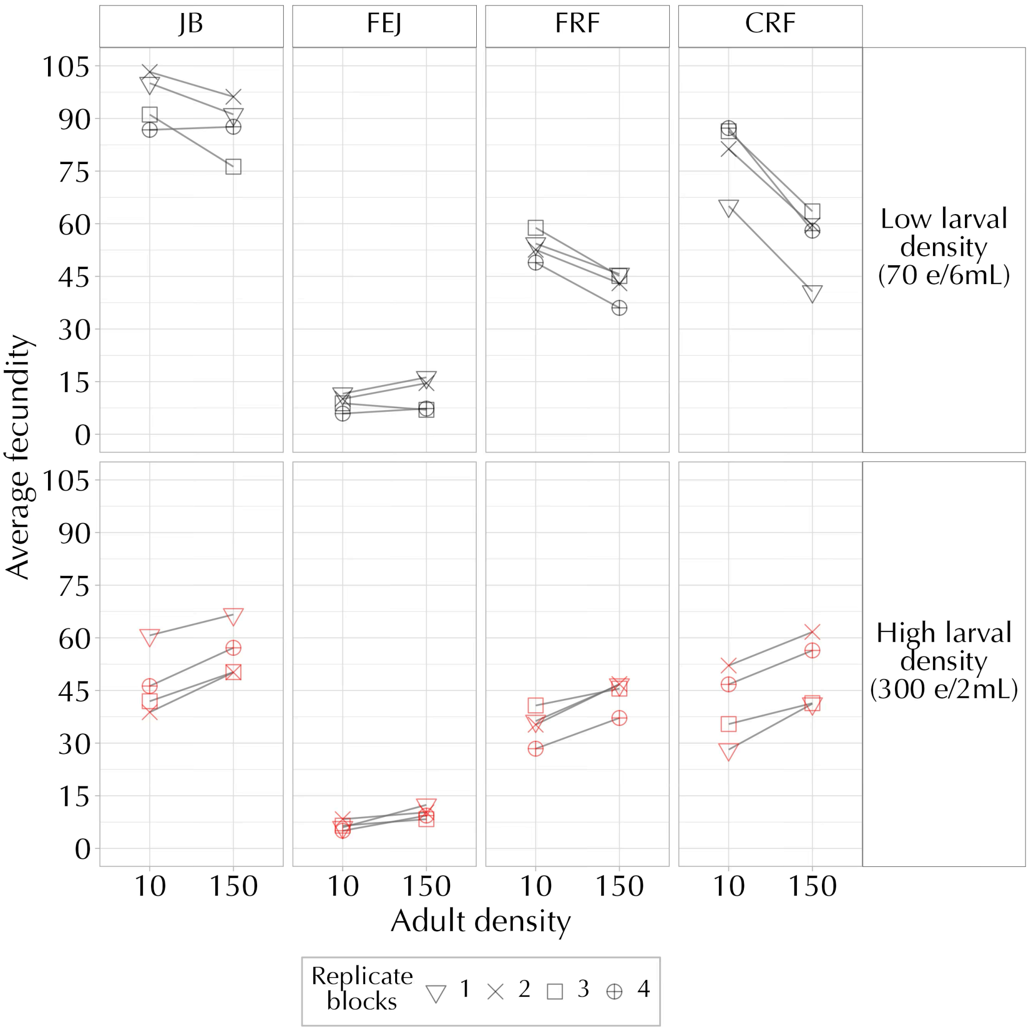


**Figure S2:** Mean female fecundity for each replicate population for all treatment combinations of selection regime, larval and adult density (averaged over day of measurement). Our experimental design cannot test any effects that involve the replicate blocks for statistical significance.
